# Cellects: a software to quantify cell expansion and motion

**DOI:** 10.1101/2024.03.26.586795

**Authors:** Aurèle Boussard, Manuel Petit, Patrick Arrufat, Audrey Dussutour, Alfonso Pérez-Escudero

## Abstract

Cellects is a user-friendly and open-source software for automated quantification of biological growth, motion, and morphology from 2D image data and time-lapse sequences (2D + t), acquired under a wide range of experimental conditions and biological systems (from fungal colonies to unicellular branching networks). The software is available as a stand-alone version, featuring a graphical interface that supports interactive parameter tuning, visualization, validation, and batch processing. The analysis pipeline can be extended and customized using a dedicated Python API.

The typical inputs and outputs are as follows. Cellects is designed to process grayscale or color images originating from standard microscopy, macroscopic imaging setups, or camera-based platforms. The software supports single or multiple organisms growing or moving in one or several arenas and can analyze multiple folders sequentially. All quantitative results (area, circularity, orientation axes, centroid trajectories, oscillations, network topology…) are exported as standardized .csv files suitable for downstream statistical analysis, ensuring reproducibility and integration into existing workflows.

## Statement of need

Modern imaging generates high-resolution biological datasets across scales, yet automated analysis remains challenging for non-experts, necessitating accessible tools.

Cellects is suited to biological systems exhibiting continuous growth, deformation, or collective motion, such as fungal colonies (Figure 1a-d), HeLa cells (Figure 1e-h), and slime molds (Figure 1i-n). By contrast, most existing tools target single species (mainly yeast or bacteria) and fail to generalize to heterogeneous morphologies such as branching slime mold networks or collective cellular movement during proliferation.

**Figure 1.**
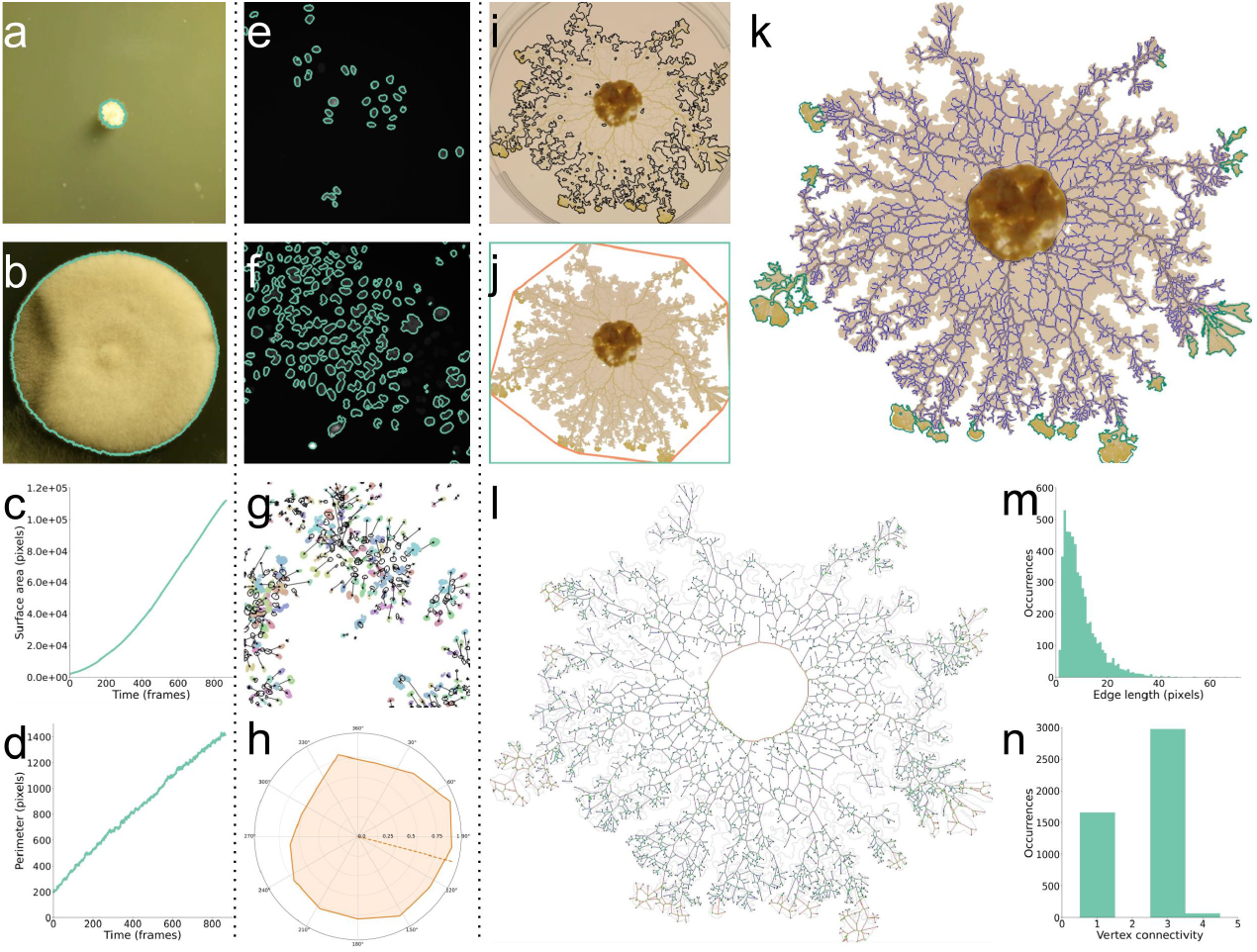
Cellular dynamics and morphologies across systems. **a,b**) Fungal growth (unknown sp.) from initial (a) to final (b) stages with green segmentation contours, from (Peñil Cobo et al., 2018); **c,d**) Corresponding area and perimeter curves over time (c: area, d: perimeter). **e,f**) Tracking of HeLa “Kyoto” cells marked with mCherry-H2B, from (Guiet, 2022) showing initial (e) and final (f) images with segmentation contours. **g**) Motion vectors (arrows) of the 250 most mobile cells among 1319 detected (black contour: original positions, colored-filled patches: final locations). **h**) Spider plot representing HeLa cell movement directions. **i,j**) *Physarum polycephalum* morphology after 16:40 hours of exploration (this study). (h: cell segmentation, i: convex hull (orange) and bounding rectangle (green)). **k,l**) Network segmentation (j: blue network; turquoise pseudopods) and graph reconstruction (k: edges colored by width from blue to red, green branching vertices, black tips, yellow food vertices). **m,n**) Physarum connectivity metrics: edge lengths (l) and vertex degrees (m). Panels a–d (fungus), e–h (HeLa), i–n (*Physarum*).

Open source alternatives often lack graphical user interfaces (GUIs) and robust automation under variable lighting/contrast conditions, while commercial platforms often require preprocessing or post-analysis using additional software, compromising reproducibility.

By combining dynamic segmentation algorithms with a modular pipeline (see Software Design), Cellects supports both single-specimen analysis and high-throughput multi-arena experiments, out-putting standardized metrics directly usable in downstream statistical workflows. While enabling reproducible studies across diverse biological models, this automated quantification reduces observer bias.

## State of the field

Cellects fills three major gaps in existing tools:

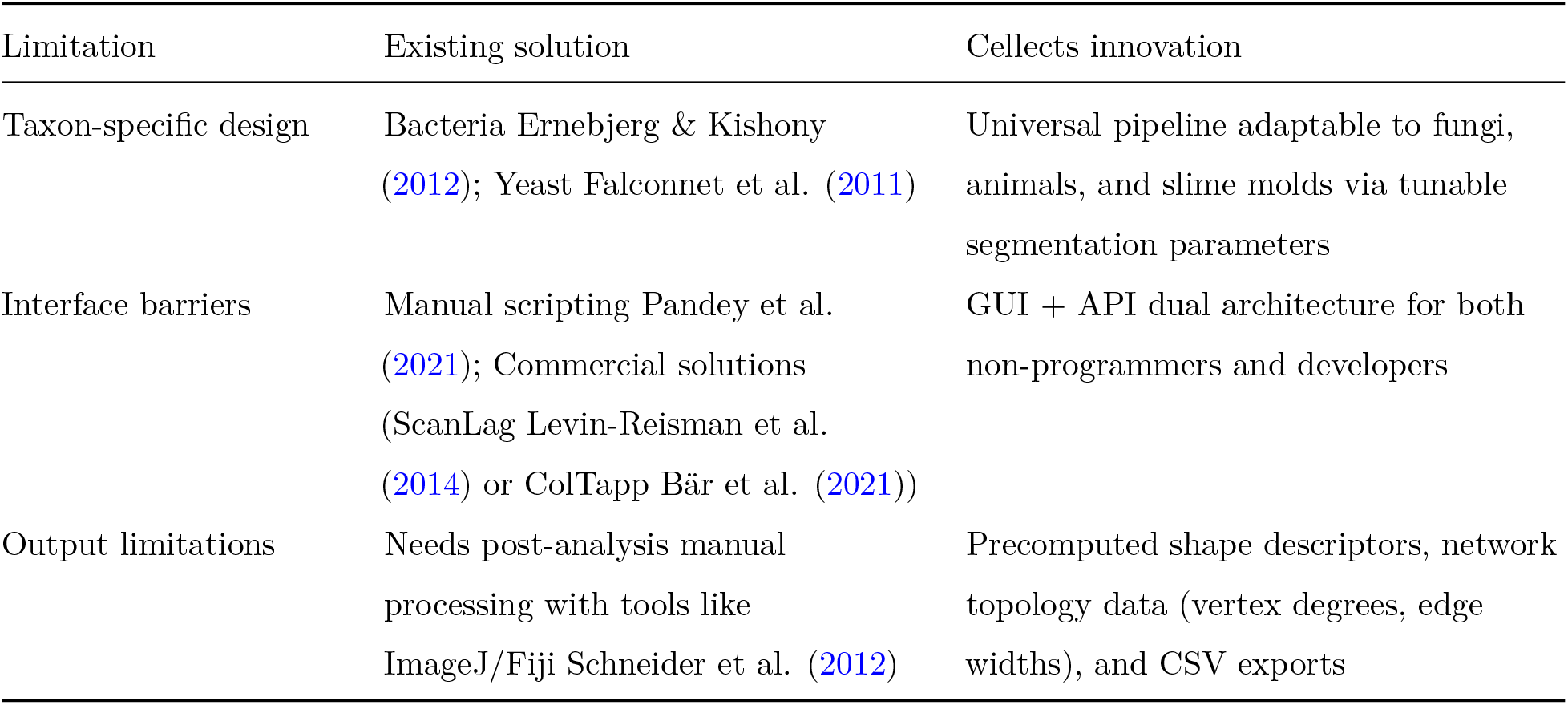

While commercial tools prioritize ease of use over customization, open-source platforms often require manual scripting. Cellects bridges this divide by combining automation with a validation tool for result refinement, enabling robust and accessible analysis of growth dynamics across biological systems and fostering reproducibility and cross-disciplinary research.

## Software design

The software is organized around a layered architecture centered on a global controller (*ProgramOrganizer*), which maintains experiment state, configuration parameters, and processing context. This controller can be driven either through the graphical user interface or programmatically.

The graphical interface follows a sequential workflow implemented via a stacked widget (*Cellects-MainWidget*), exposing successive stages for data loading (*FirstWindow*), segmentation and arena definition (*ImageAnalysisWindow*), and time-series execution (*VideoAnalysisWindow*). This structure mirrors the experimental pipeline and limits user interaction to valid analysis states while allowing iterative refinement at each step. To balance accessibility with flexibility, novice users benefit from an automated parameter search during initial setup (in ImageAnalysisWindow), while experienced ones can bypass default algorithms in advanced mode to directly customize image processing settings.

Static and temporal analyses are separated through two dedicated classes: *OneImageAnalysis*, responsible for preprocessing tasks such as greyscale conversion, filtering and background subtraction, and *MotionAnalysis*, which performs segmentation, post-processing, video-based measurements and temporal feature extraction. This separation allows computationally intensive motion analysis to build upon validated segmentation results.

Cellects targets diverse biological datasets (e.g., Fungi, HeLa cells, Myxamoebae) acquired under variable imaging conditions. Rather than relying on a single segmentation model, the image analysis layer provides automated and configurable pipelines combining multiple color-space representations, image filtering, K-means clustering and threshold-based methods. Geometric descriptors (area, perimeter, circularity) are encapsulated in the *ShapeDescriptors* class, while additional modules support graph-based dynamics, oscillatory behavior, and morphological operations.

To maintain interactivity during heavy computation, Cellects combines Qt-based threading for GUI responsiveness with multiprocessing for video analysis. Memory usage is explicitly managed through sequential image processing and controlled data release.

## Research impact statement

### Related work

Cellects’ lineage traces directly to (Vogel et al., 2015)’s MATLAB implementation, with an early iteration already employed by (Boussard et al., 2019). While the final version has not yet enabled new studies, it has been developed to address specific research questions from (Boussard et al., 2021) and from an ongoing ANR project (ANR-24-CE45-3362, PI Claire David).

### Validation

The software’s robustness is demonstrated through specimen and background accuracies (Figure 2) in diverse contexts. First, manual segmentation of Figure 1 examples provided ground truth for canonical cases: single fungus on color-varying background Figure 2a-c, multi-specimen tracking Figure 2d-f with microscopy data, and network extraction Figure 2g-i. Second, Cellects was tested on highly heterogeneous cells where manual distinction is infeasible. Accuracy was estimated via error annotations (Figure 2j) comparing original images to results, achieving >97% accuracy across five challenging conditions (Figure 2k: high contrast + optimal setup, heterogeneous colors, low contrast + desiccation, low contrast + optimal, very low resolution) through iterative parameter refinement in the GUI.

**Figure 2.**
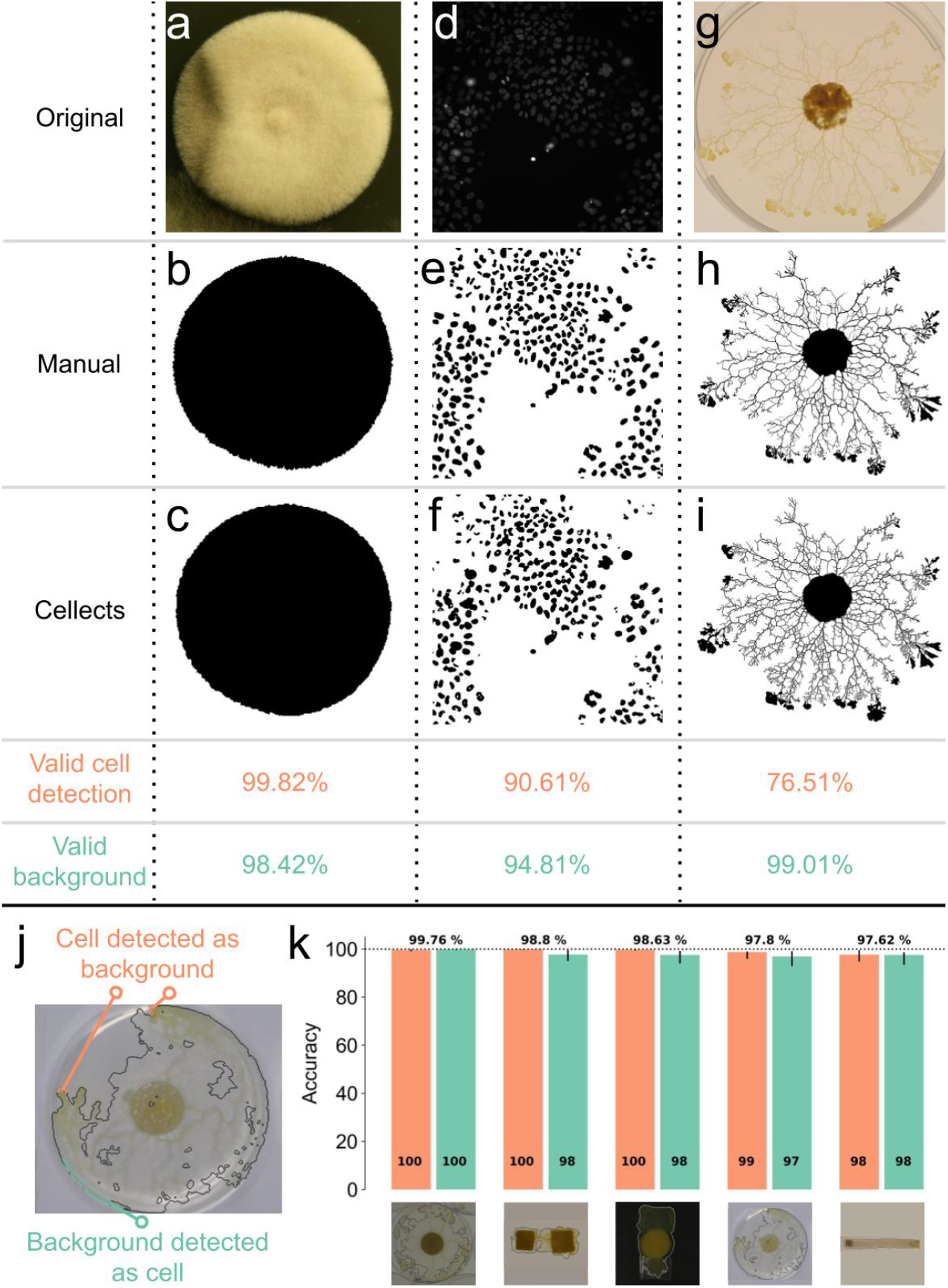
Validation of Cellects across five experimental conditions. **a,b,c**) Segmentation accuracy of the fungi (Peñil Cobo et al., 2018) against ground truth: original image (a) was used to create a mask manually (b) and using Cellects (c). The valid cell detection is the percentage of pixels accurately labelled as cells by Cellects. The valid background is the percentage of pixels accurately labelled as background by Cellects. **d,e,f**) Segmentation accuracy of the Hela “Kyoto” cells (Guiet, 2022) against ground truth. **g,h,i**) Segmentation accuracy of the *P. polycephalum* network against ground truth (this study). **j**) Image of a *P. polycephalum* plasmodia, showing the two types of errors detected during validation: background pixels classified as cell (orange), cell pixels classified as background (green). **k**) A posteriori accuracy in 5 experimental conditions (shown below the bars): high contrast with optimal setup, high contrast with heterogeneous colors, low contrast with a setup prone to desiccation, low contrast with optimal setup, low resolution. Orange: Proportion of cell pixels correctly identified as cell. Green: Proportion of background pixels correctly identified as background. Error bars show the 95% confidence interval. Percentages on top show the average of both bars.

Applications span fungal growth tracking (Figure 1a-d, (Peñil Cobo et al., 2018)), HeLa cells motion (Figure 1e-h, (Guiet, 2022)), and network analysis of *P. polycephalum* (Figure 1i-n, this study), highlighting adaptability to diverse biological systems and imaging setups.

## Acknowledgements

We thank Audrey Bizet for her work on the first experiment of Figure 2e, Charlotte Dupont and Paul-Antoine Badon for the second experiment of Figure 2e, Nirosha Murugan for her help with the fourth experiment of Figure 2e, Ana Lucía Morán Hernández for her help with the fifth experiment of Figure 2e, and Florent Le Moël, Rémi Giorno dit Journo, Guillaume Cerutti, Jonathan Legrand, and Olivier Ali for their help during software development.

## Author contribution

Conceptualization, A.B., P.A., A.D., A.P.E.; methodology, A.B., M.P., A.P.E, P.A.; software, A.B.; investigation, A.B., A.D.; resources, A.B., P.A., A.D., A.P.E.; writing – original draft preparation, A.B.; writing – review and editing, A.B., A.P.E., M.P., A.D.; visualization, A.B.; supervision, P.A., A.D., A.P.E.; project administration, A.B.; funding acquisition, A.D., A.P.E.

## AI usage disclosure

The generative model ‘Devstral-Small-2507’ was used to generate initial drafts of function docstrings and propose unit test templates. All AI-generated content underwent manual verification to ensure alignment with function usages in the context of real datasets.

## Data availability

### Lead contact

Aurèle Boussard,

### Data and code availability

The Windows and macOS versions are accessible via the following link: https://github.com/Aurele-B/Cellects/releases.

The software documentation is available at https://aurele-b.github.io/Cellects and its source code can be found at https://github.com/Aurele-B/Cellects.

To access the data and replication code, refer to: https://datadryad.org/stash/share/nCvWIZoZ8-Wnxm0CjnPbbznUPw90RYdo1YVJEQkfLIY

